# Regenerating forests around Taï National Park sustain biodiversity despite constrained floristic recovery

**DOI:** 10.64898/2026.08.11.744329

**Authors:** Aya Diane Larissa Houphouët, Yao Charles Sangne, Abdoulaye Diarrassouba, Evans Ehouman, Bruno Hérault

## Abstract

Extensive deforestation and disturbance have reshaped forest landscapes in West Africa, including areas surrounding Taï National Park, the largest remaining block of Upper Guinean rainforest. Although many degraded areas are currently undergoing regeneration, the capacity of these secondary forests to recover their biodiversity attributes remains insufficiently assessed within this park. This study quantifies the multidimensional restoration of biodiversity across 125 secondary forest plots and 15 primary forest plots representing 14,731 inventoried individuals in four sectors of the park. Using a Bayesian modelling framework, we estimate recovery trajectories of Shannon diversity, floristic composition, functional traits (wood density, leaf mass per area, seed mass), and conservation-relevant species. The different biodiversity attributes recovered at contrasting rates. Diversity was restored more rapidly (λ = 0.06) than floristic composition (λ = 0.03), and these recovery varied according to environmental variables. Among these, the presence of remnant trees showed the highest median on diversity (0.53 ± 0.61) and floristic composition (0.37 ± 0.17), promoting the rapid recovery of both attributes, followed by prior land use, particularly cocoa farming which also positively influenced the recovery of alpha diversity (λ = 0.03) and floristic composition (λ = 0.02). Regarding functional traits, they displayed contrasting dynamics: specific leaf area and seed mass recovered rapidly along successional gradients, whereas wood density followed a more gradual recovery trajectory. From a conservation perspective, although the proportion of IUCN Red List species remained stable along the successional gradient, old-growth indicator species were significantly more abundant in primary forests. To ensure the long-term conservation of biodiversity, protecting both primary and regenerating forests is essential to preserve the ecological resilience of this park.

## Introduction

Tropical forests in West Africa harbour exceptionally high levels of biodiversity and form part of one of the world’s 25 biodiversity hotspots [1,2]. Despite their ecological importance, these forests have experienced intense deforestation in recent decades, resulting in substantial biodiversity loss [3]. In Côte d’Ivoire, forest conversion has been primarily driven by agricultural expansion, especially cocoa, coffee, and oil palm alongside logging, charcoal production, infrastructure development, and mining [3–5]. These transformations have led not only to biodiversity decline but also to the degradation of ecosystem services and climate regulation.

Against this backdrop of widespread forest loss, restoring degraded lands has emerged as an essential complementary strategy to the conservation of remaining old-growth forests [6]. Secondary forests now represent a substantial proportion of tropical forest cover [7–9] and may contribute meaningfully to biodiversity conservation [10,11]. However, recovery is complex and does not progress equally across ecological features. Although species diversity often increases relatively rapidly following agricultural abandonment [12,13], floristic composition may take decades to converge toward old-growth conditions [14,15]. Moreover, recovery trajectories are strongly shaped by land-use intensity, soil degradation, remnant trees that enhance seed dispersal, and by the connectivity between neighboring landscapes [16–19].

Importantly, biodiversity recovery cannot be fully understood from taxonomic patterns alone. Beyond changes in species richness and composition, ecological succession entails shifts in species’ life-history strategies and functional attributes [20]. Early successional stages are typically dominated by fast-growing pioneer species, whereas late-successional and shade-tolerant species establish progressively as forest structure develops [14,21]. These successional dynamics are reflected in changes in functional traits, which provide mechanistic insight into ecosystem functioning and resilience [22]. Traits such as wood density, leaf mass per area (LMA), and seed mass are closely linked to growth–mortality trade-offs, resource-use strategies, and regeneration processes [23–26]. Seed mass, in particular, mediates the balance between colonization ability and establishment success [27–29]. Integrating functional composition with taxonomic and compositional analyses therefore provides a more comprehensive framework for evaluating forest recovery.

In West Africa, where remaining old-growth forests are extremely limited and deforestation rates remain among the highest globally, conservation increasingly depends on the capacity of secondary forests to sustain biodiversity. Yet it remains unclear whether regenerating forests can meaningfully substitute for primary forests in maintaining endangered species, functional diversity, and ecosystem processes. Addressing this uncertainty is critical for guiding restoration strategies, landscape planning, and protected-area buffering. These issues are especially critical in landscapes subjected to intense anthropogenic pressure, such as those surrounding Taï National Park in south-western Côte d’Ivoire. Taï National Park constitutes the largest remaining continuous block of Upper Guinean humid rainforest and one of the last extensive remnants of tropical rainforest in West Africa. As a major stronghold of regional biodiversity, it harbours numerous endemic and threatened species. However, episodes of degradation particularly during the 2002-2011 politico-military crises have affected both the park and its periphery. Since 2015, degraded areas have been undergoing natural regeneration [30], creating a mosaic of intact old-growth forest, disturbed old-growth forest, and secondary forests of varying ages. In such a globally significant forest landscape, understanding the extent and pace of biodiversity recovery is essential for informing regional conservation and restoration strategies.

To date, few studies in Upper Guinean forests have jointly quantified taxonomic, functional, compositional, and conservation-relevant recovery using reference-based models. Here, we provide one of the first multidimensional assessments of natural forest regeneration surrounding Taï National Park, integrating community composition, functional traits, biomass, and species conservation status within a unified Bayesian framework. To provide conceptual clarity, we define “recovery” as the degree of convergence of biodiversity attributes in secondary forests toward reference conditions observed in intact old-growth forests. Recovery therefore refers not only to increases in species diversity over time, but also to compositional similarity and functional convergence relative to old-growth benchmarks. This reference-based approach allows us to distinguish between rapid gains in taxonomic diversity and slower or incomplete recovery of compositional and functional attributes. Building on this framework, we analyse biodiversity recovery trajectories in secondary forests surrounding Taï National Park. Specifically, we address the following questions: (i) What are the recovery trajectories of Shannon diversity and floristic composition, and which factors influence them? (ii) How do key functional traits such as wood density, leaf mass per area, and seed mass change along secondary succession, and can they serve as indicators of forest recovery? and (iii) Do secondary forests contribute to the persistence and recovery of conservation-relevant species?

## Methods

### Study Area

The study was conducted in and around Taï National Park (TNP), located in south-western of Côte d’Ivoire between 5°15′ to 6°07′ N and 7°25′ to 7°54′ W (Fig 1). TNP constitutes the largest remaining continuous block of Upper Guinean humid rainforest and is recognised as a UNESCO World Heritage Site [31,32]. The park covers approximately 5,360 km² and represents one of the last extensive remnants of primary tropical rainforest in West Africa [33]. The region is characterised by a humid tropical climate with two rainy seasons (March to July and September to November) and two relatively drier periods. Mean annual rainfall ranges between 1,700 and 2,200 mm, and mean annual temperature is approximately 25–27 °C. Soils are predominantly ferralitic and vary locally depending on topography and drainage conditions. Vegetation is dominated by evergreen and semi-deciduous moist forest. The park harbours high levels of plant and vertebrate diversity, including numerous endemic and threatened species characteristic of the Upper Guinean forest block [34,35]. Surrounding landscapes have been extensively transformed by agricultural expansion, particularly cocoa cultivation, as well as logging and artisanal gold mining.

**Fig. 1.**
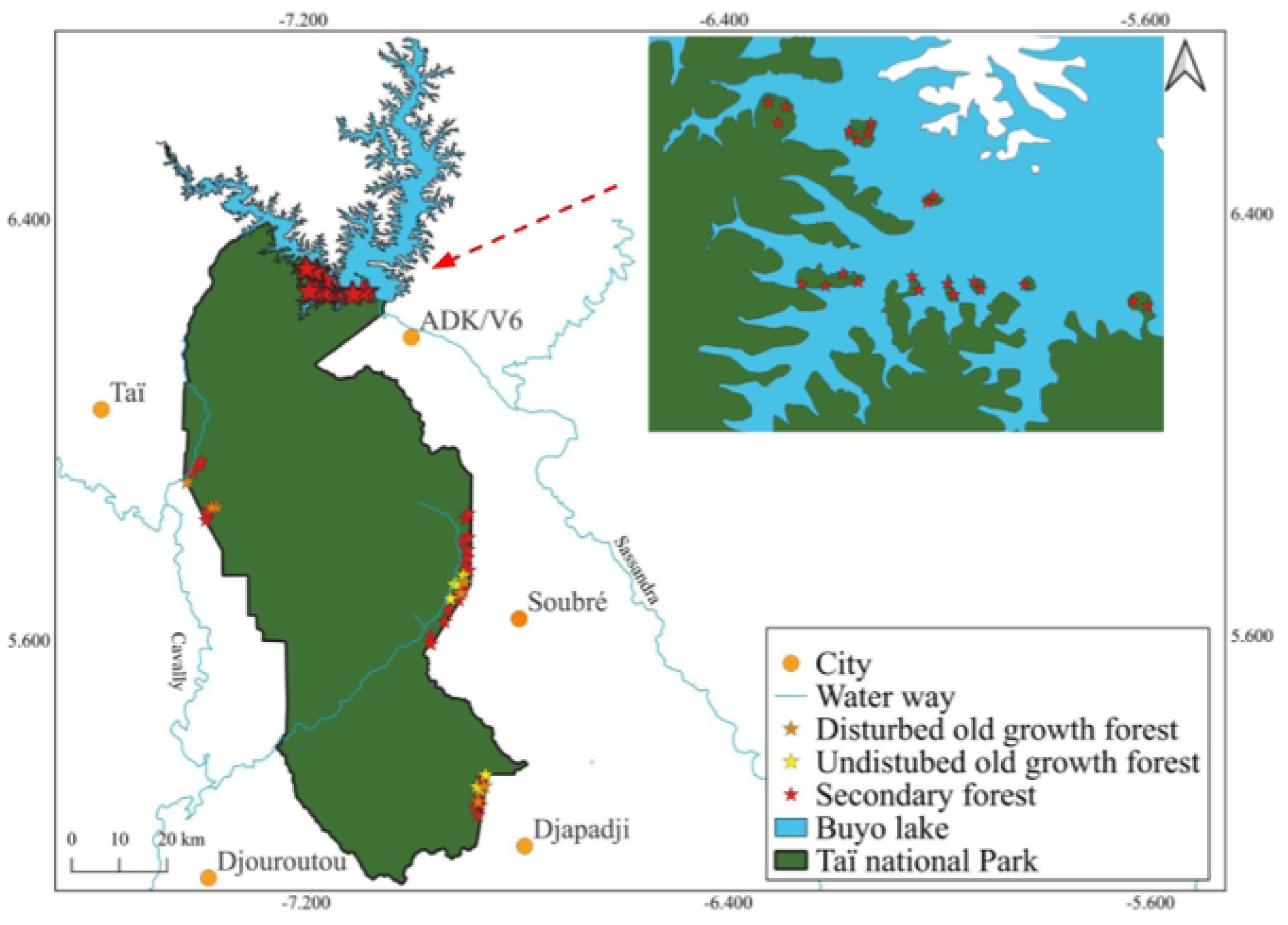
Spatial distribution of sampling plots across Taï National Park and surrounding landscapes in south-western Côte d’Ivoire. Sampling plots represent a chronosequence of secondary forests (9–26 years since abandonment; n = 125) and reference intact old-growth forests (n = 15). Plots are distributed within and around park boundaries. The map also shows Buyo Lake, major waterways, and nearby cities to provide geographic context. The inset highlights the northern sampling cluster near Buyo Lake, where forest fragmentation has created a mosaic of secondary and old-growth stands. Printed under a CC BY license, with permission from OIPR, original copyright [2026].

During the post-electoral crises of 2000 and 2010, reduced surveillance led to forest encroachment and degradation in parts of the park and its periphery. Since 2015, restoration and natural regeneration processes have been ongoing in previously degraded areas [36]. As a result, the current landscape forms a mosaic of intact old-growth forest, disturbed old-growth forest, and secondary forests of varying ages originating from former agricultural lands or mining sites.

### Vegetation Data

A total of 140 vegetation plots were established in and around Taï National Park. Of these, 125 plots were located in secondary forests and 15 plots in intact old-growth forests, which served as reference sites. Each plot measured 25 × 25 m (0.0625 ha). Secondary forests represented a chronosequence ranging from 9 to 26 years since abandonment. Forest age was determined based on information from local communities, park management records, and field evidence of past land use. Following the post-electoral crisis period, agricultural activities inside the park were discontinued and cocoa plantations were progressively removed as part of restoration and management interventions. Subsequent natural succession gave rise to stands of varying ages, forming the basis of the chronosequence analysed in this study. Within each 25 × 25 m plot, all woody stems with a diameter at breast height (DBH) ≥ 2.5 cm were inventoried. DBH was measured at 1.3 m above ground level. For multi-stemmed individuals, each stem meeting the diameter threshold was recorded separately. Lianas were excluded from the inventory. Species were identified in the field by experienced botanists. When identification was uncertain, specimens were examined using the regional floras of Aké-Assi [37] and Hutchinson & Dalziel [38] to ensure accurate determination. Although voucher specimens were not systematically deposited in a public herbarium, taxonomic verification procedures were applied to ensure consistency and reliability across plots. Nomenclature was standardised according to the Angiosperm Phylogeny Group IV (APG IV) classification system, and species names were harmonised to avoid synonymy or spelling inconsistencies within the dataset.

All original data used in this study will be publicly available and can be accessed from Zenodo upon acceptance. Permissions for this study were granted by the Office Ivoirien des Parcs et Réserves (OIPR), the governmental body responsible for managing Taï National Park, Côte d’Ivoire.

### Functional Trait Data

We analysed three functional traits: wood density (WD), leaf mass per area (LMA), and seed mass. These traits were selected because of their established links to growth–mortality trade-offs, resource-use strategies, regeneration processes, and successional dynamics in tropical forests [23,24].

Wood density values were retrieved using the R package *BIOMASS* [39] through the function *getWoodDensity()*, which assigns species-level values when available and substitutes genus-level averages when species-specific data are missing. This procedure ensured complete coverage of wood density values across all species in the dataset. Seed mass and LMA values were obtained primarily from the TRY global plant trait database [40], including regional datasets integrated into TRY [41]. Seed mass data were additionally compiled from the SER-SID database (ser-sid.org). For species lacking values in these databases, seed mass measurements were supplemented using published data from [42].

When multiple trait values were available for a given species across datasets, the median value was used to reduce the influence of extreme observations. Species-level trait values were then assigned to all corresponding individuals in the vegetation dataset. Trait coverage varied among traits. Wood density values were available for 100% of species, seed mass for approximately 49% of individuals, and LMA for 56% of individuals.

### Environmental variables

To account for ecological and anthropogenic drivers of recovery, we recorded a set of environmental and disturbance-related variables expected to influence regeneration processes (Table 1).

- Large and medium-sized mammals were included because they play a key role in seed dispersal and recruitment dynamics in tropical forests. Their presence can enhance colonization by late-successional species and influence community composition. Mammal occurrence was assessed through indirect field signs of species known to function as seed dispersers or large herbivores.
- Remnant trees were quantified as indicators of biological legacies from pre-disturbance forests. Such trees can provide local seed sources, structural complexity, and perching sites for frugivores, thereby potentially accelerating succession.
- Soil hydromorphy was recorded as a proxy for edaphic constraints on species establishment and growth, as waterlogged conditions can limit recruitment of certain functional groups.
- Island configuration was included to capture landscape isolation effects, which may reduce propagule influx and alter recovery trajectories.
- Signs of ongoing human activities were documented to account for continued disturbance pressure that may slow or redirect successional dynamics.

**Table 1.** Plot-level environmental variables used to assess biotic legacies, edaphic constraints, landscape isolation, and ongoing anthropogenic activities potentially influencing biodiversity recovery in secondary forests surrounding Taï National Park, Côte d’Ivoire.

| Variables | Description | Values |
| --- | --- | --- |
| Mammal | Number of large/medium-sized mammal species detected through indirect signs | 0-3 |
| Hydromorphy | Presence of hydromorphic soil features in surface horizons | 0/1 |
| Island | Plot located in isolated island | 0/1 |
| Remnants | Density of isolated old-growth trees persisting after land conversion | trees.ha<sup>-1</sup> |
| Activities | Number of disturbance types observed (poaching, agriculture,mining) | 0-3 |

Together, these variables capture biotic, abiotic, and anthropogenic drivers of succession and were incorporated as covariates in subsequent models of biodiversity recovery.

### Conservation-relevant species

To evaluate the contribution of secondary forests to biodiversity conservation, we considered two categories of conservation-relevant species: (i) old-growth indicator species and (ii) species listed under threatened categories of the IUCN Red List.

Indicator species characteristic of old-growth forests were identified using the method of [43], implemented in the R package *indicspecies*. We compared two groups of plots: intact old-growth forests and secondary forests younger than 20 years. The indicator value (IndVal) combines measures of specificity and fidelity to a given habitat type, allowing the identification of species strongly associated with old-growth conditions. Species were considered significant old-growth indicators when their indicator value was associated with a permutation test P-value < 0.002. A stringent significance threshold was applied to reduce the risk of false positives and to ensure robust identification of species reliably associated with old-growth forest conditions. For each plot, we subsequently quantified the occurrence and abundance of these indicator species.

For all inventoried species, conservation status was assessed using the IUCN Red List of Threatened Species (https://www.iucnredlist.org/). Species classified as Vulnerable (VU) or Endangered (EN) were considered conservation-relevant for this analysis. The presence and abundance of these species were quantified across forest types to evaluate their persistence and potential recovery during secondary succession.

### Biodiversity metrics

Diversity was quantified using Shannon diversity, expressed as the Hill number of order 1 (q = 1). Diversity of order 1 corresponds to the exponential of Shannon entropy and represents the effective number of species in a community. Estimates were computed using the R package *entropart* [44]. Sample coverage was first estimated for each plot using the function *coverage()*. As coverage was consistently above 0.90 across plots, diversity was estimated at a standardized coverage level of 0.90 to ensure comparability among plots. This coverage-based standardization minimizes bias due to incomplete sampling while avoiding distortions caused by unequal sampling effort.

Composition reflects both species identity and their relative abundances within each plot. To quantify compositional similarity between secondary forests (SF) and intact old-growth forests (OGF), we calculated the abundance-based Chao–Jaccard similarity index for all possible SF– OGF plot pairs. This index accounts for unseen shared species and corrects for undersampling, thereby providing a robust estimate of compositional similarity. To define the reference state, similarities among old-growth plots (OGF–OGF) were also calculated. These within-OGF similarities represent the natural variability of intact forests and define the benchmark toward which secondary forests are expected to converge during succession [15]. Although the 15 reference plots are modest relative to the park’s total area, they were distributed across four sectors of the park to capture part of its spatial heterogeneity, and their pairwise similarity was used to explicitly characterise — rather than assume — the natural variability of old-growth forest composition used as the recovery benchmark.

### Modeling Strategies

All models were built within a Bayesian framework. A Bayesian approach was chosen because of its flexibility in handling complex hierarchical structures, its ability to accommodate different likelihood distributions across response variables, and its explicit quantification of uncertainty through posterior distributions. A log-normal likelihood was used because all response variables considered (diversity, composition, and functional traits) are strictly defined on the positive real line (ℝ⁺) and exhibited right-skewed distributions [45]. Log-normal likelihoods were implemented in Stan using its standard parameterization, with mean parameters defined on the original scale and passed to the distribution through the log link. Models were implemented in Stan and fitted using Hamiltonian Monte Carlo (HMC) sampling for parameter estimation [46].

### Q1-Biodiversity Recovery Trajectories

To estimate recovery trajectories and intrinsic recovery rates, we applied the exponential recovery model developed by [47]. This model assumes that the instantaneous recovery rate of an ecosystem attribute is proportional to the distance between its current state and its asymptotic value [45]. Such a formulation captures saturating recovery dynamics commonly observed during secondary succession.

For secondary forests, the likelihood model for a biodiversity attribute *X* (diversity or composition) in plot *p* at time *t* since abandonment was specified as:

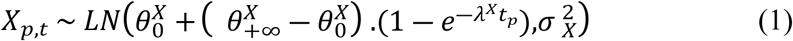

where:

- *X_p_*_,*t*_ is the observed value of attribute *X* in plot *p* at time *t*,
- θ^X^_0_ represents the initial value of *X* at the beginning of the recovery process,
- θ^X^_+∞_ is the asymptotic value of *X*, corresponding to the long-term equilibrium conditions,
- ë*^X^* is the intrinsic recovery rate controlling the speed of convergence toward the asymptote,
- σ^2^_X_ represents the residual variance among secondary forest plots.

For intact old-growth forests, time since regeneration is not defined and attribute values are assumed to represent the asymptotic state. The likelihood was therefore specified as:

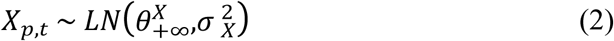

This formulation allows old-growth forests to directly inform the asymptotic parameter θ^X^_+∞_.

To account for anthropogenic disturbance effects in old-growth forests, asymptotic values were adjusted by including a disturbance effect term:

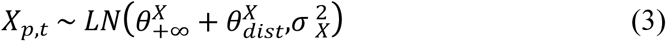

where θ^X^_dist_ represents the effect of disturbance intensity on the asymptotic value of attribute *X*.

To evaluate how environmental and disturbance-related variables influenced recovery dynamics, covariates (Table 1) were incorporated into the recovery rate parameter λ^X^_p_. Because recovery rates are strictly positive, we modelled them using an exponential link function:

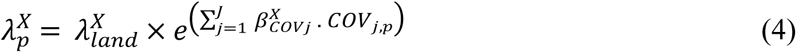

where:

- λ^X^_land_ is the baseline recovery rate associated with previous land use,
- β^X^_COVj_ represents the effect of covariate *j* on recovery rate,
- *COV_j_*_,*p*_ is the standardized value of covariate *j* for plot *p*.

All continuous covariates were standardized prior to analysis to facilitate parameter interpretation and comparison of effect sizes.

### Q2 - Functional trait dynamics

To analyse how functional traits change during succession, we used an individual-level Bayesian model linking trait values to both stem diameter *DBH* and forest age *t*. This approach captures shifts in trait distributions along the tree size gradient during recovery, consistent with the idea that secondary forests progressively approach old-growth trait–size relationships as forests mature [47]. Because trait values were strictly positive and right-skewed, they were modelled using a log-normal likelihood. For each tree *n* in secondary forests, trait values *Y* (wood density, leaf mass per area, or seed mass) were modelled as:

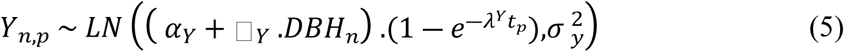

where *DBH_n_* is the diameter of individual *n*, *t_p_* is time since abandonment for the plot *p* where the individual occurs, *α_Y_* is the baseline value of the trait when the diameter effect is null, ◻*_Y_* quantifies the effect of DBH on the expected trait value, *λ^Y^* is the convergence rate toward the asymptotic state, and σ^2^_y_ is the residual variance. Under this formulation, the expected trait–size relationship converges toward the *α_Y_* + _◻_*_Y_. DBH_n_* as forest age increases, thereby allowing trait distributions to progressively align with old-growth conditions along the DBH gradient.

For intact old-growth forests, trait values were assumed to reflect the asymptotic (reference) trait–size relationship:

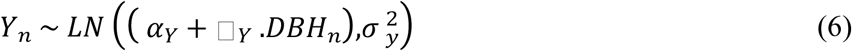

To account for possible anthropogenic disturbance shifts away from intact reference conditions, we included an additive disturbance effect on the expected trait–size relationship:

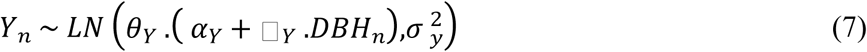

where *θ_Y_* represents the effect of disturbance intensity on the expected value of trait *Y* (i.e., *θ_Y_* > 1 indicates higher expected trait values under disturbance, whereas *θ_Y_* < 1 indicates lower expected trait values).

### Q3 - Conservation-relevant species

To assess whether conservation-relevant species are maintained or recovering during succession, we calculated, for each plot, the proportion of individuals belonging to (i) old-growth indicator species and (ii) IUCN-listed species (Vulnerable or Endangered). Secondary forests were grouped into two age classes (≤ 11 years and > 11 years since abandonment) and compared with intact old-growth forests. This comparison allowed us to evaluate whether the relative representation of conservation-relevant species increases with forest age and approaches old-growth conditions.

## Results

Across all forest types, we recorded 14,731 woody individuals (DBH ≥ 2.5 cm), corresponding to 174 species distributed among 138 genera and 48 families.

### Biodiversity recovery trajectories

Shannon diversity and floristic composition did not recover at the same rate (Fig 2, Table 2). Shannon diversity increased rapidly during succession, reaching approximately 75% of its asymptotic value within the first 40 years. The intrinsic recovery rate was λ = 0.06 (90% CI: 0.02–0.17). Diversity started at an estimated 8.94 effective species (θ₀) and converged toward an asymptotic value of 20.39 in old-growth forests (θh∞). In contrast, floristic composition recovered more slowly (λ = 0.03; 90% CI: 0.00–0.16). Because λ is constrained to be strictly positive by the model’s exponential link function, this interval does not literally include zero; however, its lower bound is very close to zero, indicating that a near-negligible, practically stagnant recovery rate cannot be excluded. On average, secondary forests required over 40 years to approach half of the asymptotic similarity observed in old-growth forests (Fig 2). Composition increased from an initial value of 0.19 to an asymptote of 0.72 (Table 2). Previous land use influenced recovery rates. Recovery was faster in former cocoa farms than in former mining sites for diversity, whereas land-use effects on compositional recovery were negligible (Table 2).

**Fig. 2.**
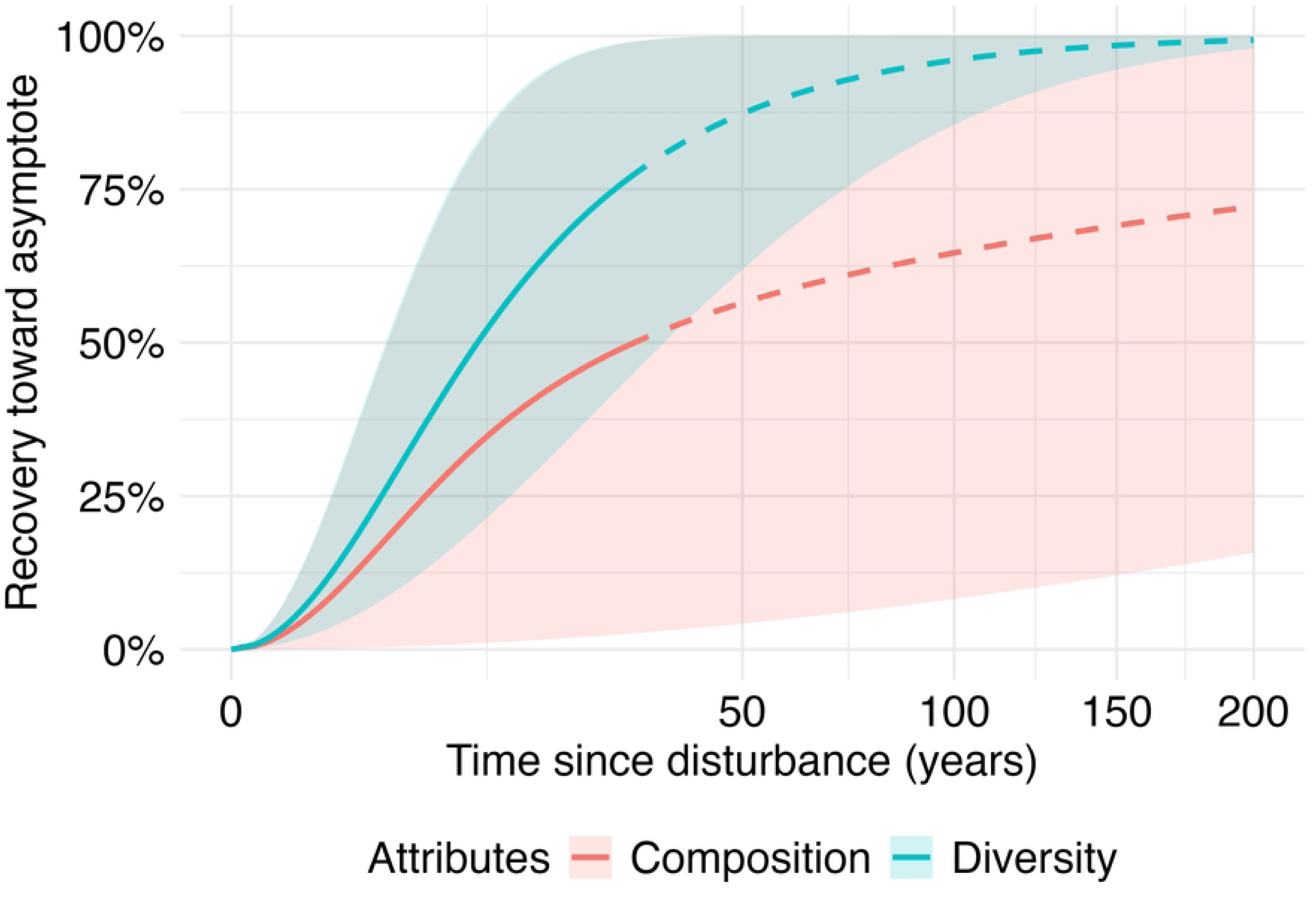
Modelled recovery trajectories of Shannon diversity and floristic composition in secondary tropical forests of West Africa (Taï National Park, Côte d’Ivoire). Lines represent posterior mean recovery trajectories estimated from the Bayesian exponential recovery model; shaded areas indicate 95% credible intervals. Solid lines correspond to the temporal range robustly informed by the underlying 9–26 year chronosequence (up to ∼40 years since disturbance); dashed lines represent the model’s extrapolation beyond this range, up to 200 years, and should be interpreted with caution given the increasing uncertainty. Recovery is expressed as the percentage of convergence toward old-growth reference conditions, with 100% corresponding to the asymptotic state observed in intact West African old-growth forests.

**Table 2.**
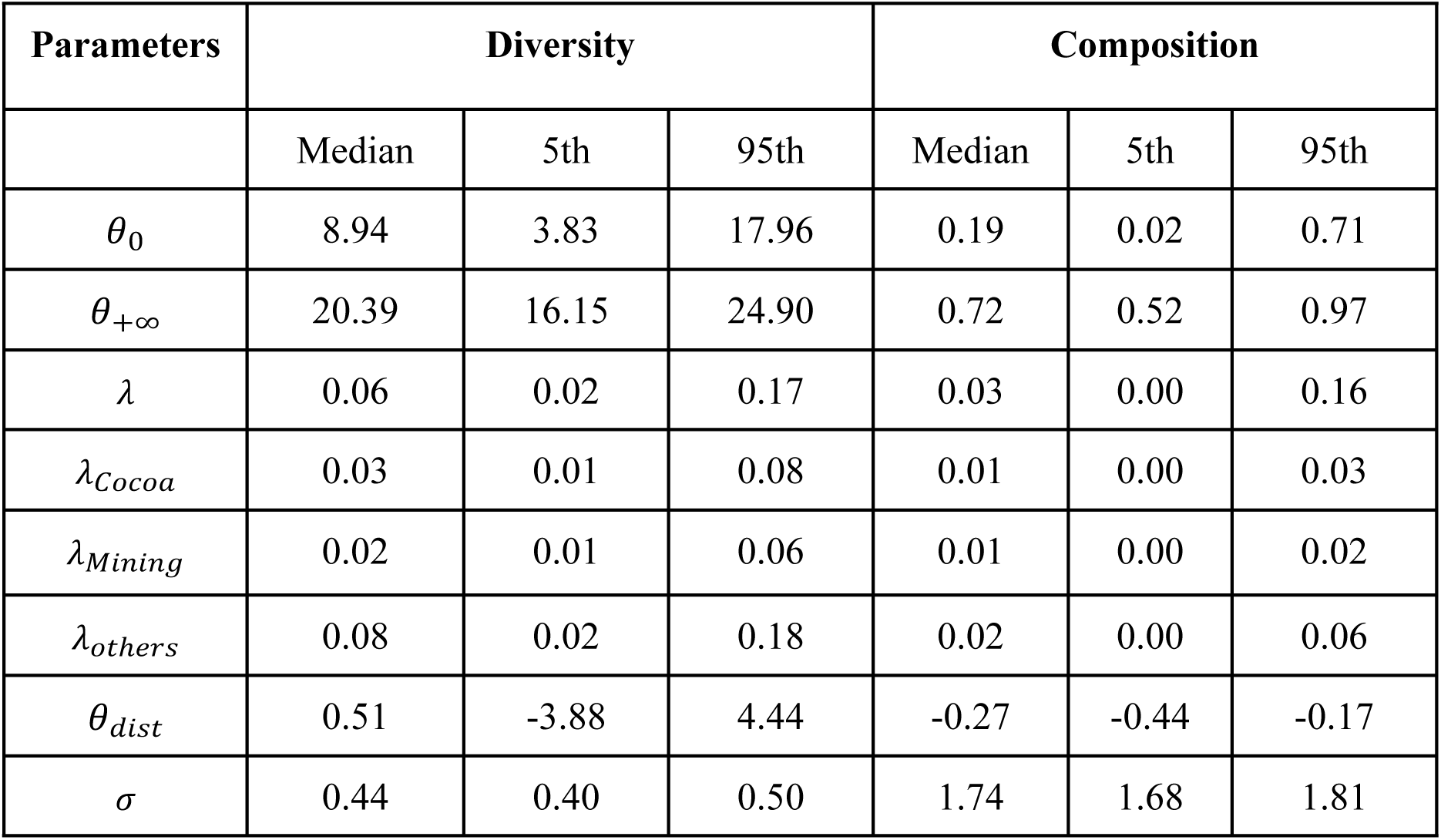
Posterior summaries of the recovery model parameters for diversity and composition in secondary forests of West Africa. Posterior medians and 90% credible intervals (5th–95th percentiles) of the exponential recovery model parameters for Shannon diversity (Hill number of order 1) and floristic composition (abundance-based Chao–Jaccard similarity). *θ*_0_ represents the initial value of the recovery trajectory; *θ*_+∞_ the asymptotic value corresponding to old-growth reference conditions; *λ* the intrinsic recovery rates; *θ_dist_* the effect of past disturbance on the asymptotic value; and σ the residual variance of the log-normal likelihood. Posterior estimates of environmental covariate effects (β parameters) are not shown here and are presented in Fig. 3.

Environmental covariates had contrasting effects on recovery rates (Fig 3). Remnant trees showed positive effects on both diversity and composition. Human activities and island configuration were associated with slower recovery of both attributes. Mammal presence increased compositional recovery but was negatively associated with diversity recovery. Soil hydromorphy and disturbance exhibited variable effects, with wide credible intervals for diversity but negative effects on compositional recovery.

**Fig. 3.**
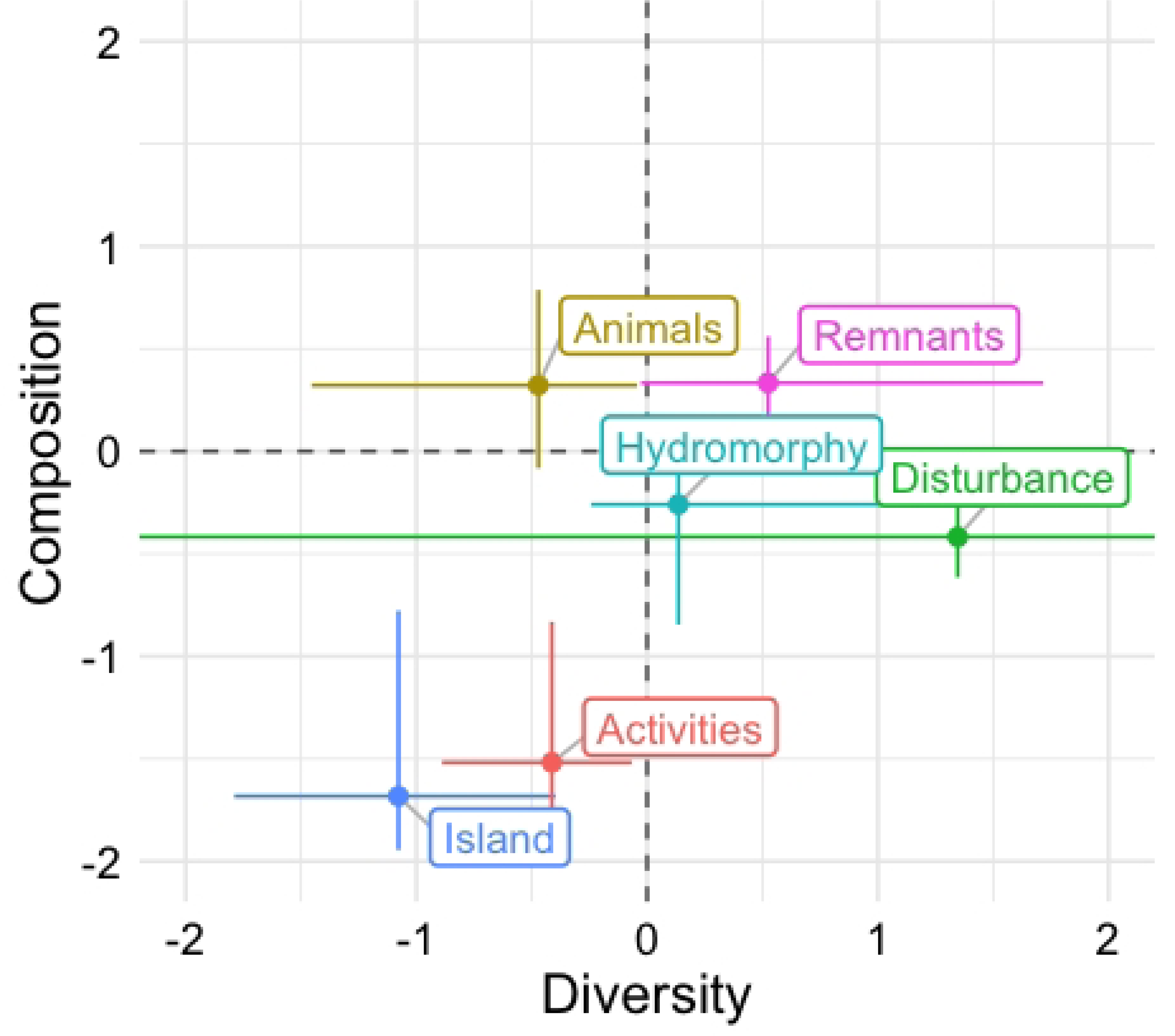
Environmental drivers of diversity and composition recovery rates. Points represent posterior mean effects of standardized covariates on the intrinsic recovery rate (λ) for diversity (x-axis) and composition (y-axis). Error bars indicate 95% credible intervals. Vertical and horizontal dashed lines represent zero effect. Positive values indicate an increase in recovery rate, whereas negative values indicate slower recovery. Covariates include mammal presence, remnant tree density, soil hydromorphy, island location, human activities, and disturbance intensity.

### Functional Trait Dynamics

Trait values varied jointly with stem diameter and forest age, with distinct convergence rates among traits (Table 3).

**Table 3.** Posterior summaries of the recovery model parameters for functional traits in secondary forests of West Africa. Posterior medians and 90% credible intervals (5th–95th percentiles) of the individual-level exponential convergence model parameters for seed mass, wood density, and leaf mass per area. α represents the expected trait value when DBH = 0; β quantifies the effect of stem diameter on trait values; λ is the intrinsic convergence rate toward old-growth reference conditions; θ represents the multiplicative effect of disturbance on trait values in old-growth forests; and σ is the residual variance of the log-normal likelihood.

| Parameters | Seed mass |  |  | Wood density |  |  | Leaf mass per area |  |  |
| --- | --- | --- | --- | --- | --- | --- | --- | --- | --- |
|  | Median | 5th | 95th | Median | 5th | 95th | Median | 5th | 95th |
| $\alpha$ | 20.43 | 19.89 | 21.01 | 0.61 | 0.61 | 0.62 | 60.12 | 59.63 | 60.60 |
| $\beta$ | 1.26 | 0.41 | 2.11 | -0.17 | -0.18 | -0.16 | 4.63 | 4.63 | 5.46 |
| $\lambda$ | 1.21 | 0.72 | 2.25 | 0.15 | 0.15 | 0.16 | 1.4 | 0.9 | 2.47 |
| $\theta$ | 0.98 | 0.95 | 0.99 | 0.97 | 0.96 | 0.99 | 0.98 | 0.99 | 0.99 |
| $\sigma$ | 2.85 | 2.81 | 2.89 | 0.22 | 0.22 | 0.23 | 3.62 | 3.5 | 3.69 |

Wood density showed a gradual increase with forest age and remained structured by stem diameter (Fig 4). Larger individuals exhibited slightly lower wood density values than smaller individuals, consistent with the negative size effect (β = −0.17). Convergence toward old-growth conditions was slow (λ = 0.15). The disturbance parameter (θ = 0.97) indicated slightly lower expected values under disturbance relative to intact old-growth forests.

**Fig. 4.**
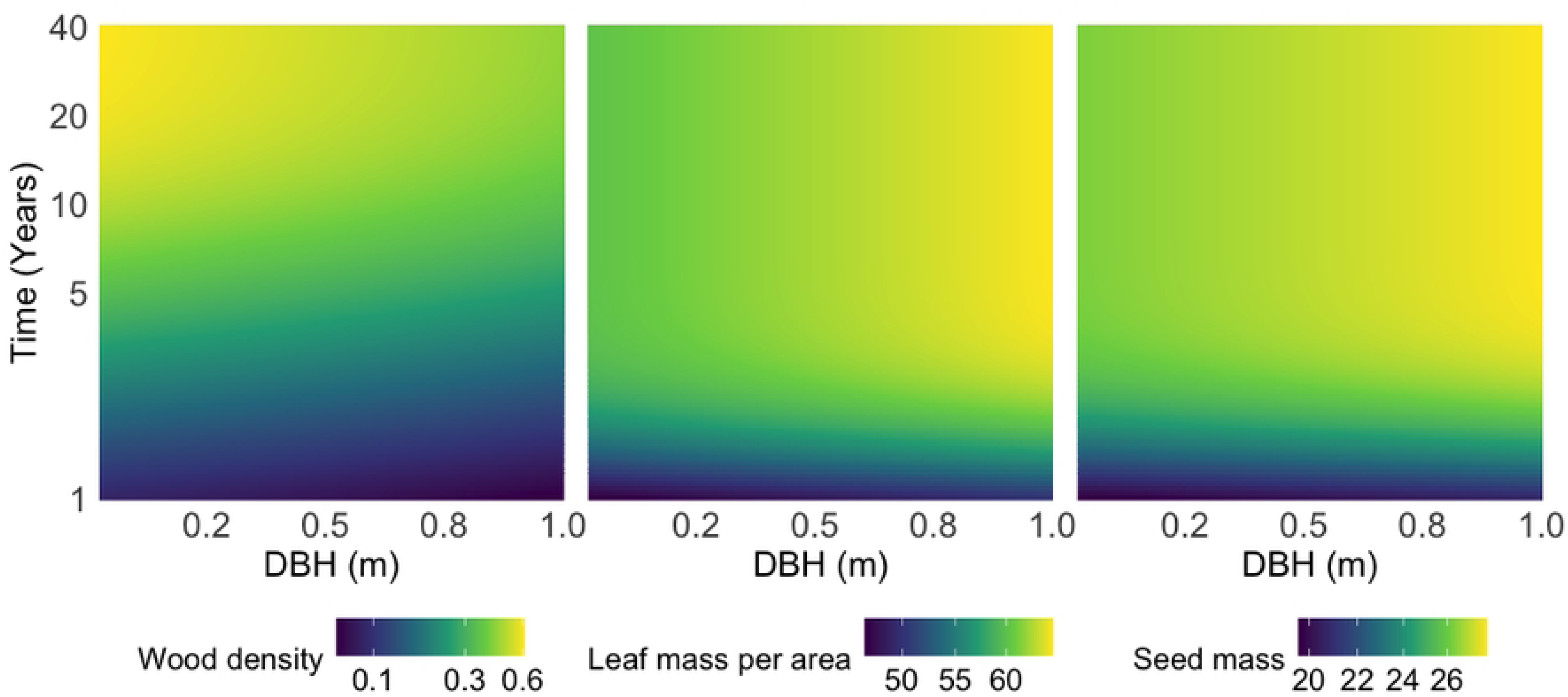
Modelled trajectories of functional trait distributions along the DBH and successional age gradients in secondary forests of West Africa. Heatmaps represent posterior mean predicted values of wood density (g.cm^-3^), leaf mass per area (LMA; g.m^-2^), and seed mass (mg) as functions of stem diameter (DBH, m; x-axis) and time since abandonment (years; y-axis). Color gradients indicate expected trait values derived from the individual-level convergence model. Predictions illustrate progressive alignment of trait–size relationships toward old-growth reference conditions as forest age increases.

LMA was strongly influenced by stem diameter, with larger trees displaying higher values (β = 4.63). In contrast to wood density, LMA converged rapidly toward old-growth reference conditions (λ = 1.40). Predicted values exceeded 60 g.m^-2^ within the first decade of succession across most of the DBH gradient (Fig 4). The disturbance effect was minimal (θ = 0.98).

Seed mass increased with stem diameter (β = 1.26) and exhibited rapid convergence dynamics (λ = 1.21). Predicted seed mass values for larger individuals exceeded 25 mg within the first 10 years of succession and remained stable thereafter (Fig 4). The disturbance parameter was close to unity (θ = 0.98; 90% CI: 0.95–0.99), indicating limited deviation from intact old-growth reference values.

### Conservation-relevant species

Among the 174 recorded species, 16 (9.2%) were classified as either Endangered (EN) or Vulnerable (VU) according to the IUCN Red List (Appendix 1), and 44 species (25.3%) were identified as old-growth indicator species based on the IndVal analysis (Appendix 2).

IUCN-listed species were present across all forest stages. Their proportion remained relatively stable among young secondary forests (≤ 11 years), older secondary forests (> 11 years), and intact old-growth forests, with values ranging between 8% and 9% of individuals (Fig 5).

**Fig. 5.**
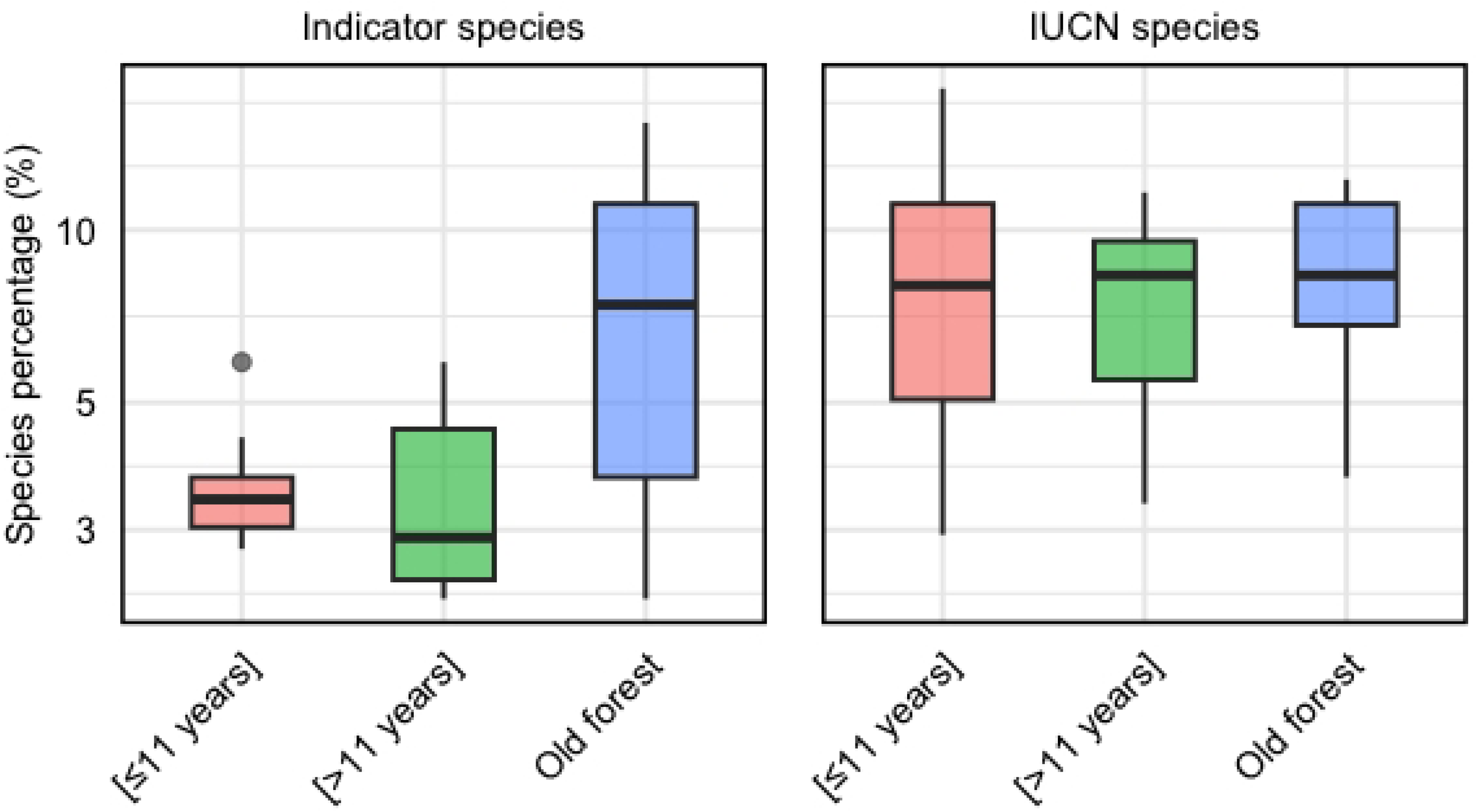
Proportion of conservation-relevant species across successional stages in West African tropical forests. Boxplots show the percentage of individuals belonging to old-growth indicator species (left panel) and IUCN-listed species categorized as Vulnerable or Endangered (right panel) across three forest categories: secondary forests ≤ 11 years since abandonment, secondary forests > 11 years, and intact old-growth forests. Boxes represent interquartile ranges, horizontal lines indicate medians, and whiskers show the range of observed values

In contrast, the proportion of old-growth indicator species increased with forest age. Indicator species represented approximately 3–5% of individuals in young secondary forests, 4–6% in intermediate secondary forests, and 7–11% in old-growth forests (Fig 5).

## Discussion

In the last extensive block of Upper Guinean rainforest in West Africa, our results show that secondary forests surrounding Taï National Park retain substantial ecological potential. Although biodiversity dimensions recover at different rates, species diversity rebounds relatively quickly, functional attributes re-establish within decades, and conservation-relevant species persist across successional stages. These findings provide encouraging evidence that, when protected from further disturbance, naturally regenerating forests in this region can contribute meaningfully to biodiversity conservation and landscape resilience.

### Rapid diversity recovery masks slower and context-dependent compositional convergence

The rapid recovery of diversity likely reflects the resilience of secondary forests and the progressive accumulation of species following land abandonment [48]. Similar patterns have been reported in Neotropical forests, where up to 90% of species richness can recover within two decades [48–50]. In contrast, floristic composition depends on the return of late-successional species, many of which are dispersal-limited and characterized by larger seed mass [51,52]. As a result, compositional convergence may require several decades to centuries. Comparable trajectories have been observed in the Congo Basin [53], the Neotropics [54], and Southeast Asia [52], reinforcing the generality of this asynchronous recovery pattern.

Importantly, compositional recovery was not only slower but also highly variable, reflecting strong context dependence. It should be noted that the lower bound of the credible interval for the compositional recovery rate is very close to zero, meaning our data cannot exclude a near-negligible recovery rate that would be practically equivalent to stagnation over the sampled chronosequence (9–26 years); this uncertainty highlights the need for longer-term monitoring to distinguish slow recovery from an effective stall. Previous land use played a central role. Abandoned cocoa plantations often retain partial soil fertility [55] and remnant trees [56,57], which facilitate regeneration and canopy re-establishment [58,59]. In contrast, artisanal gold mining removes topsoil, disrupts seed banks, alters soil structure, and may lead to heavy metal accumulation, reducing microbial activity and long-term fertility [16,60]. Such disturbances strongly constrain recolonization by forest specialists and slow compositional convergence [41].

Environmental factors further modulated recovery trajectories. Remnant trees emerged as a key driver accelerating both diversity and compositional recovery. These trees create favorable microhabitats, increase structural heterogeneity, and act as local seed sources [55,61]. They also attract seed dispersers such as birds and bats, facilitating the recruitment of late-successional species [62,63]. Similar patterns have been documented elsewhere, where species richness and compositional similarity to primary forests increase around remnant trees after two decades of succession [64,65]. Conversely, human activities and forest isolation slowed recovery. Hunting and logging associated with gold mining reduce populations of large seed dispersers, thereby limiting recruitment of large-seeded forest species [66,67]. Forests on islands created a fragmentation experience that reduced propagule influx and strong edge effects, favoring pioneer and generalist species [68–70]. In line with island biogeography theory [71], limited connectivity constrains long-term compositional recovery. Hydromorphic soils also exerted contrasting effects, promoting herbaceous dominance (e.g., Marantaceae) that can inhibit woody recruitment and slow structural and compositional reassembly [59,72–74].

Together, these results show that while diversity rebounds relatively quickly in regenerating forests of West Africa, floristic composition follows a slower and highly context-dependent trajectory. Around Taï National Park, this asynchrony underscores the importance of maintaining landscape connectivity, remnant trees, and effective protection from continued disturbance to support long-term compositional recovery.

### Rapid reassembly of leaf and regeneration traits contrasts with slow structural maturation

Functional traits recovered at markedly different rates, revealing distinct ecological dimensions of succession in the forests surrounding Taï National Park. In intact old-growth forests, trait distributions reflect conservative resource-use strategies adapted to closed-canopy conditions, where light is limiting and competition for water and nutrients is intense [75]. Dominant species tend to be shade tolerant and invest in durable tissues, including dense wood and structurally robust leaves.

In contrast, secondary forests initially host species that prioritize rapid growth over structural investment. The lower wood density observed in younger forests likely reflects the dominance of pioneer species that colonized soon after abandonment. These heliophilous species capitalize on high light availability created by disturbance, investing in fast height growth rather than dense, mechanically resistant wood. As succession proceeds and canopy closure increases, later-arriving species face reduced light and heightened resource competition [76]. These species typically allocate more biomass to dense wood, enhancing mechanical stability and resistance to disturbance [77]. Such ontogenetic and successional shifts in species composition likely explain the slow convergence of wood density, consistent with broader evidence that structural maturation in tropical forests requires several decades [21,78].

In contrast, leaf mass per area (LMA) reassembled rapidly. LMA increased with both forest age and stem diameter, reflecting adjustments to vertical light gradients and hydraulic constraints. Leaves positioned higher in the canopy experience greater irradiance and wind exposure, promoting thicker and denser leaf construction [79,80]. Increased hydraulic path length in taller individuals further constrains water transport to the canopy, favoring structural reinforcement of leaves. Smaller trees and understory individuals, by contrast, operate under lower light intensity and shorter hydraulic distances, allowing thinner and lighter leaves. The rapid convergence of LMA suggests that canopy stratification and associated microclimatic gradients re-establish relatively quickly during succession. Because LMA is assigned as a fixed species-level value to every individual of a given species, rather than measured directly on individual trees, our data cannot capture in situ or ontogenetic adjustment of leaf traits within individuals. The observed convergence in LMA therefore necessarily reflects progressive species turnover toward taxa with characteristically higher LMA, not physiological or plastic adaptation of existing trees. In addition, LMA values were available for only 57% of individuals, and species lacking a trait record were excluded from this analysis; while we have no indication of systematic bias in which species were missing, this limited coverage should be kept in mind when assessing the generality of the LMA convergence pattern.

Seed mass dynamics reflect a distinct regeneration axis. Pioneer species typically produce numerous small and light seeds that disperse effectively into disturbed environments [29,81]. Although these seeds contain limited reserves, they benefit from high-light conditions and rapid germination. In contrast, shade-tolerant species produce fewer but larger seeds rich in reserves, enhancing seedling survival under low-light and competitive understory conditions [82–84]. Small-seeded species compensate for lower individual survival through high fecundity and persistence in soil seed banks [81,85]. The relatively rapid stabilization of seed mass distributions indicates that regeneration strategies diversify early in succession, even before full structural maturation of the forest.

Together, these patterns show that while leaf-level adjustments and regeneration strategies reassemble within decades, structural attributes associated with long-lived woody tissues recover more slowly. Around Taï National Park, this decoupling between rapid functional adjustment and delayed structural maturation highlights that forest recovery involves multiple ecological dimensions operating on different temporal scales.

### Naturally regenerating forests contribute meaningfully to conservation

The presence of IUCN-listed species across all forest stages indicates that naturally regenerating forests around Taï National Park maintain conservation value throughout succession. The proportion of Endangered and Vulnerable species remained relatively stable across young secondary, older secondary, and old-growth forests, suggesting that early successional habitats contribute to sustaining threatened taxa. Similar complementary roles of secondary forests have been reported in other tropical landscapes [86–88].

In contrast, old-growth indicator species were markedly more abundant in intact forests than in secondary stands (Fig. 5). Although indicator species were present in both young and older secondary forests likely reflecting the persistence of remnant trees [89], their proportion remained substantially lower than in old-growth forests. This pattern suggests that while secondary forests support elements of primary forest composition, full recovery of old-growth identity requires longer time spans and sustained protection. Comparable trends have been documented in West Africa, where indicator species progressively increase but remain less abundant in secondary stands than in primary forests [73].

Together, these findings highlight the complementary but not substitutive role of secondary forests in the Upper Guinean region. While they do not yet replicate the full compositional integrity of old-growth forests, their capacity to maintain threatened species and retain primary-forest elements underscores their importance for biodiversity conservation in West Africa.

## Conclusion

Following episodes of intense degradation during the post-electoral crises, the forests surrounding Taï National Park have undergone a decade of natural regeneration. Our results show that this regeneration is not merely structural [59,90] but ecologically meaningful. Secondary forests recover species diversity rapidly, reassemble key functional attributes within decades, and retain threatened taxa across successional stages. Although full compositional convergence toward old-growth forests remains slower and strongly context-dependent, the progressive return of primary-forest elements demonstrates substantial ecological resilience in the last major block of Upper Guinean rainforest. In a region where deforestation has drastically reduced forest cover, these findings provide concrete evidence that protecting regenerating forests around Taï is not a secondary objective but a central pillar of long-term conservation. Safeguarding both intact old-growth stands and naturally recovering forests is essential to maintain biodiversity, ecological functions, and landscape connectivity in West Africa.

## Acknowledgments

The authors wish to thank the management of Taï National Park for their logistical and financial support, as well as the forest rangers whose assistance greatly facilitated the data collection phase

## Supporting information

**Appendix S1. IUCN-listed tree species recorded in Taï National Park (Côte d’Ivoire).** Species classified as Endangered (EN) or Vulnerable (VU) according to the IUCN Red List. Values of 1 indicate the assigned conservation category

**Appendix S2. Old-growth indicator species identified using the Dufrêne–Legendre indicator value method.** Indicator species associated with intact old-growth forests, identified using the Dufrêne–Legendre indicator value (IndVal) method. Only species with significant association (P < 0.002) are reported. IndVal values range from 0 (no association) to 1 (perfect specificity and fidelity to old-growth forests).

